# Chrysin, but not flavone backbone, decreases anxiety-like behavior in animal screens

**DOI:** 10.1101/575514

**Authors:** León Jesús German-Ponciano, Bruna Patrícia Dutra Costa, Leonardo Miranda Feitosa, Kimberly dos Santos Campos, Suianny Nayara da Silva Chaves, Jonathan Cueto-Escobedo, Monica Lima-Maximino, Juan Francisco Rodríguez-Landa, Caio Maximino

## Abstract

Chrysin (5,7-dihydroxyflavone), a nutraceutical flavonoid present in diverse plants, has a backbone structure shared with the flavone backbone, with additional hydroxyl groups that confers its antioxidant properties and effects at the GABA_A_ receptor complex. However, whether these effects are due to the hydroxyl groups is unknown. Here we report the effects of chrysin or the flavone backbone (1 mg/kg) in rats subjected to the elevated plus-maze and the locomotor activity test, as well as in the zebrafish evaluated in light/dark model. Chrysin, but not flavone, increased entries and time in the open arms of the elevated plus-maze, as well as time on white compartment of the light/dark model in zebrafish. These effects were comparable to diazepam, and were devoid of motor effects in both tests, as well as in the locomotor activity test. On the other hand, flavone decreased risk assessment in the light/dark test but increased rearing in the locomotor activity test in rats, suggesting effects threat information gathering; important species differences suggest new avenues of research. It is suggested that the specific effects of chrysin in relation to flavone include more of a mechanism of action in which in addition to its action at the GABA_A_/benzodiazepine receptor complex also could be involved its free radical scavenging abilities, which require specific research.

**Preprint:** https://doi.org/10.1101/575514;

**Data and scripts:** https://github.com/lanec-unifesspa/chrysin

## Introduction

Flavonoids are polyphenolic compounds present in diverse plants (Ghosh & Scheepens, 2009; Krafczyk et al., 2009) and have been considered as nutraceuticals due to they are part of the human diet (Marventano et al., 2017) and produce diverse pharmaceuticals actions. Also, they are able to cross the blood-brain barrier and modify the brain function (Johnston et al., 2015). Particularly, the flavonoid chrysin (5,7-dihydroxyflavone) has been studied by its antioxidant properties (Siddiqui et al., 2018), but currently also due to its neuropharmacological actions produced on specific brain structures implicated in neuropsychiatric disorders (Filho et al., 2016; Bortolotto et al., 2018). Chrysin is a natural flavonoid present in honey, propolis and diverse plant extracts (German-Ponciano et al., 2018), and is one of the most studied polyphenolic nutraceuticals (Chadha et al., 2017). Chrysin is used as a nutraceutical as a “testosterone boosting agent” (a claim that is probably very exaggerated; Gambelunghe et al., 2003), but there is some preclinical evidence that this molecule ameliorates indices of hepatic and renal functioning in diabetic animals (Ramanathan & Thekkumalai, 2014; Yamamoto, 2014). There is also preclinical evidence that this flavonoid exerts anxiolytic-like effects (Wolfman et al., 1994; Salgueiro et al., 1997; Zanoli et al., 2000; Rodríguez-Landa et al., 2019).

Chrysin has a backbone structure that consists in two aromatic rings (A and C), and a phenyl B ring, which is attached to the second position of ring C and shares the flavone backbone, with an additional hydroxyl group in the 5 and 7 positions of the A ring (Mani et al., 2017; Figure 1). The potential of chrysin to act as free radical scavenger has been attributed to the presence of these hydroxyl groups (Sathiavelu et al., 2009; Souza et al., 2015), and it has been suggested that this functional group represents the main action site of this flavonoid to produce a great variety of potential therapeutic effects to ameliorate diverse physiological and psychiatric disorders (Sheela & Augusti, 1995). However, no studies have directly compared the effects of similar doses of chrysin and the flavone backbone on the potential therapeutic effects reported at preclinical research.

**Figure 1:**
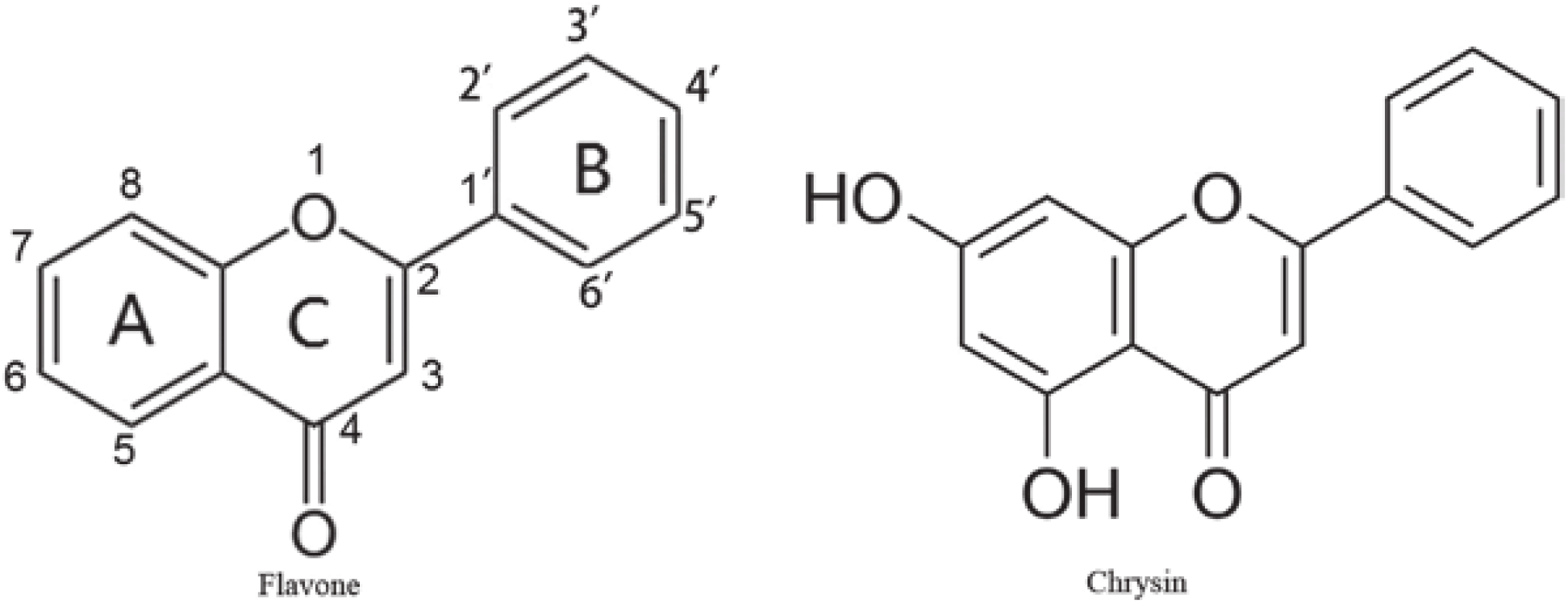
Chemical structures of flavone and chrysin.

Chrysin (either isolated from *Passiflora Coerulea, Passiflora incarnata, Matricaria Chamomilla* or as a synthetic drug) exerts anxiolytic-like effects through action on the GABA_A_ receptors without the associated side effects (i.e. sedation, amnesia, myorelaxation) that have been attributed to other GABA_A_ modulators (benzodiazepines, ethanol, etc.); these effects have been evaluated in behavioral models using rodents (Wolfman et al., 1994; Salgueiro et al., 1997; Zanoli et al., 2000; Rodríguez-Landa et al., 2019). Although it has been suggested that presence of the hydroxyl groups in the chrysin structure is the responsible for this pharmacological action, this effect has not been compared with the effects produced by the flavone backbone in preclinical models of anxiety-like behavior.

In addition to mammalian model organisms used to study the anxiolytic-like effects of diverse substances, non-mammalian models have been developed to explore the potential therapeutic effects of drugs in a short time (Norton & Bally-Cuif, 2010). The use of zebrafish in preclinical research in the psychopharmacology field has increased recently. Biobehavioral assays of anxiety and screening tests for anxiolytic-like effects, such as the rodent light/dark test (LDT), have been adapted to zebrafish, considering that behavioral phenotypes are similar between rodents and zebrafish (Maximino et al., 2010a). Behavioral screens using zebrafish have evaluated the effects of clinically effective anxiolytic drugs, including benzodiazepines (Bencan et al., 2009; Gebauer et al., 2011; Maximino et al., 2011; Schaefer et al., 2015) and ligands of both the central and peripheral BZD sites (Lima-Maximino et al., 2018), suggesting good pharmacological isomorphism and the ability of such screens to identify potential anxiolytic drugs that can progress in the drug development pipeline. In addition to increasing the toolbox of behavioral screens for psychoactive drugs, zebrafish screens and biobehavioral assays also increase our understanding of the “core” mechanisms of behavioral functions, such as fear and anxiety, that are altered in human patients and shared between fish, rodents, and humans (Gerlai, 2014).

The aim of the present study was to compare the effects of the nutraceutical flavonoid chrysin and the flavone backbone (the basic structure of the flavone without the hydroxyl radical groups), using the LDT in zebrafish, as well as the elevated plus-maze (EPM) and locomotor activity test in rats. This could contribute to identify if anxiolytic-like effects produced by chrysin are due to the presence of hydroxyl radicals in its basic structure, which could impact in the specific design of novel molecules destined to pharmacotherapy of anxiety. This manuscript is a complete report of all the studies performed to test the effect of chrysin, flavone, or diazepam on anxiety-like behavior in rats and zebrafish. We report all data exclusions (if any), all manipulations, and all measures in the study.

## Materials & methods

### Animals

#### Rats

32 adult male Wistar rats (2.5 months old), weighing 200-250 g, were used. Animals were housed in Plexiglas cages (4-5 rats per cage) in the Laboratorio de Neurofarmacología (Universidad Veracruzana, Xalapa, Mexico) under a 12 h/12 h light/ dark cycle (lights on at 7:00 AM) at 25°C ± 1°C with *ad libitum* access to food and water. The experimental procedures were performed in accordance with national (*Especificaciones Técnicas para la Producción, Cuidado y Uso de Animales de Laboratorio* NOM-062-ZOO-1999) and international (National Research Council (US) Committee for the Update of the Guide for the Care and Use of Laboratory Animals, 2011) ethical recommendations. All efforts were made to minimize animal discomfort during the study.

#### Zebrafish

48 adult unsexed zebrafish of the longfin phenotype were used in the present experiments. Animals were acquired in a local aquarium shop (at Belém, Brazil) and kept in collective tanks at the Laboratório de Neurofarmacologia e Biofísica (UEPA, Marabá, Brazil) for at least 2 weeks before experiments started. Conditions in the maintenance tank were kept stable, as described by Lawrence (2007) and recorded in protocols.io (Maximino et al., 2019; [https://dx.doi.org/10.17504/protocols.io.rupd6vn]). Recommendations in the Brazilian legislation (Diretriz brasileira para o cuidado e a utilização de animais para fins científicos e didáticos - DBCA. Anexo I. Peixes mantidos em instalações de instituições de ensino ou pesquisa científica, 2017) were followed to ensure ethical principles in animal care and throughout experiments.

### Drugs and treatments

Chrysin (CAS #480-40-0; Sigma-Aldrich, C80105) and flavone (CAS #525-82-6; Sigma-Aldrich, F2003) were dissolved in 5% dimethyl sulfoxide (DMSO) and injected *i.p.* For all drugs, the dose was 1 mg/kg, a dose of chrysin shown to be anxiolytic in elevated plus-maze (Wolfman et al., 1994). Thirty-two rats were assigned to four independent groups (*n* = 8/group). The vehicle group received 5% DMSO solution (Golden Bell Reactivos, México City, México) in which chrysin and diazepam were prepared; a second group (CHR) received 1 mg/kg chrysin; a third group (FLAV) received 1 mg/kg flavone; and a fourth group (DZP) received 2 mg/kg diazepam (CAS #439-14-5; Tocris, 2805; German-Ponciano et al., 2020) as a pharmacological control of anxiolytic activity. Sixty minutes after the injection, the rats were evaluated in the elevated plus maze and then in the locomotor activity test. The time elapsed between injection and the behavioral test has previously demonstrated anxiolytic-like effects of both chrysin and diazepam at the doses that were uesed in present study (Taiwo et al., 2012; Rodríguez-Landa et al., 2019; Cueto-Escobedo et al., 2020; German-Ponciano et al., 2020).

Zebrafish were randomly drawn from the holding tank immediately before injection and assigned to four independent groups (*n* = 12/group). Groups were identical to those used for rats, including vehicle (5% DMSO). The injection volume was 1 μL/0.1 g of body weight, and animals were injected intraperitoneally (Kinkel et al., 2010). The order with which groups were tested was randomized via generation of random numbers using the randomization tool in http://www.randomization.com/. Experimenters were blinded to treatment by using coded vials for drugs. The data analyst was blinded to phenotype by using coding to reflect treatments in the resulting datasets; after analysis, data was unblinded.

### Elevated plus-maze

The rats were separately evaluated in the elevated plus maze, and then in the locomotor activity test. On the testing day, the rats were brought to the experimental room at 11:00 AM and left for 1 h to acclimate to the novel surroundings. The behavioral tests began at 12:00 AM. The apparatus consisted of two opposite open and closed arms set in a “+” configuration, and it was situated in an illuminated room at 40 lux. The open arms were 50 cm length × 10 cm width, and the closed arms were 50 cm length × 10 cm width, with 40 cm-high walls. The entire maze was elevated 50 cm above the floor (Walf & Frye, 2007). To evaluate the effects of the treatments, the rats were individually placed in the center of the maze, facing an open arm. The following variables were scored during 5 min:

1. *Number of entries into the open arms* (N);
2. *Number of entries into the closed arms* (N);
3. *Total number of entries* (open arms + closed arms), N;
4. *Percentage of open arm entries* (Proportion of entries made in the open arms in relation to the total number of entries)
5. *Time spent on the open arms* (s);
6. *Time spent in head-dipping* (s);
7. *Number of head-dipping events* (N);
8. *Time spent in stretched-attend posture* (SAP; s);
9. *Number of SAP events* (N).

After the elevated plus maze test, the rats underwent the locomotor activity test. Approximately 2 min elapsed between tests.

### Locomotor activity test

Each rat was individually placed in the locomotor activity cage (44 cm length × 33 cm width × 20 cm height). The floor of the cage was delineated into twelve 11 cm × 11 cm squares to evaluate spontaneous locomotor activity, grooming, and rearing for 5 min. At the beginning of the test, the rat was gently placed in one of the corners of the cage. The following variables were scored:

1. *Number of squares crossed* (crossings; a crossing was considered when the rat passed from one square to another with its hind legs);
2. *Time spent rearing* (rearing was considered when the rat emitted a vertical position relative to the cage floor);
3. *Time spent grooming* (including paw licking, nose/face grooming, head washing, body grooming/scratching, leg licking, and tail/genital grooming).

Digital video cameras (Sony DCR-SR42, 40 optical zoom, Carl Zeiss lens) were installed above each apparatus (elevated plus maze and locomotor activity test) to record activity. Two independent observers measured the behavioral variables using *ex profeso* software to record the number and time (in seconds) of each evaluated behavioral variable until >95% agreement was reached among the observers. After each test session, the apparatus was carefully cleaned with a 10% ethanol solution to remove the scent of the previous rat, which can influence spontaneous behavior of the subsequent rat.

### Light/dark test

The light/dark preference (scototaxis) assay was performed as described elsewhere (Maximino, 2018 [https://10.17504/protocols.io.srfed3n]; Maximino et al., 2010b). Briefly, 30 min after injection animals were transferred individually to the central compartment of a black/white tank (15 cm height × 10 cm width × 45 cm length) for a 3-min acclimation period, after which, the doors which delimit this compartment were removed and the animal was allowed to freely explore the apparatus for 15 min. While the whole experimental tank was illuminated from above by a homogeneous light source, due to the reflectivity of the apparatus walls and floor average illumination (measured just above the water line) above the black compartment was 225 ± 64.2 (mean ± S.D.) lux, while in the white compartment it was 307 ± 96.7 lux. The following variables were recorded:

1. *Time spent on the white compartment:* the time spent in the white half of the tank (*s*);
2. *Transitions to white:* the number of entries in the white compartment made by the animal throughout the session;
3. *Entry duration:* the average duration of an entry (time on white / transitions);
4. *Erratic swimming:* defined as the number of zig-zag, fast, and unpredictable swimming behavior of short duration;
5. *Freezing:* the duration of freezing events (*s*), defined as complete cessation of movements with the exception of eye and operculum movements;
6. *Thigmotaxis:* the duration of thigmotaxic events (*s*), defined as swimming in a distance of 2 cm or less from the white compartment’s walls;
7. *Frequency of risk assessment:* defined as a fast (<1 s) entry in the white compartment followed by re-entry in the black compartment, or as a partial entry in the white compartment (i.e., the pectoral fin does not cross the midline);

A digital video camera (Samsung ES68, Carl Zeiss lens) was installed above the apparatus to record the behavioral activity of the zebrafish. Two independent observers, blinded to treatment, manually measured the behavioral variables using X-Plo-Rat 2005 (https://github.com/lanec-unifesspa/x-plo-rat). Inter-observer reliability was at least > 0.95.

### Statistical analysis

Data were analyzed using Asymptotic General Independence Tests, a permutation-based analogue of one-way ANOVA, followed by pairwise permutation tests, analogous to pairwise t-tests with *p-*values adjusted for the false discovery rate. Analyses were made in R (R Core Team, 2019), using the package ‘coin’ (Hothorn et al., 2006). Normality was not assumed, and thus no specific test for normality was performed; however, this type of analysis is resistant to deviations from the assumptions of the traditional ordinary-least-squares ANOVA, and is robust to outliers, thus being insensitive to distributional assumptions (such as normality) (Mangiafico, 2015). This test is based on the permutation of a statistic maxT, the maximum of several competing univariate statistics. Outliers were detected using an *a priori* rule based on median absolute deviation (MAD) of time on white (the main endpoint of the LDT) or time in the open arms (the main endpoint of the EPM), with values above or below 3 MADs being removed (Leys et al., 2013). Using this procedure, one outlier was removed from the flavone-treated group in zebrafish.

### Open science practices

Hypotheses and methods were not formally pre-registered. Data and analysis scripts can be found at a GitHub repository (https://github.com/lanec-unifesspa/chrysin).

## Results

### Elevated plus-maze

In the elevated plus-maze (Figure 2A), animals made more entries in the open arms (Figure 2B; maxT = 2.4407, p = 0.05) after the treatment with chrysin (p = 0.02) and diazepam (p = 0.02) in comparison to the control group. No effect was found for entries in the closed arm (Figure 2C; maxT = 1.0295, p = 0.63) or total entries (Figure 2D; maxT = 1.5331, p = 0.36). A significant effect was found for percentage of entries in the open arms (Figure 2E; maxT = 2.7848, p = 0.02), but differences disappeared after adjusting *p-*values in *post-hoc* contrasts. A significant effect was found for time spent in the open arms (Figure 2F; maxT = 2.6129, p = 0.03), this variable was increased after treatment with chrysin (p = 0.02) and diazepam (p = 0.01). No effects were found for time spent head-dipping (Figure 2G; maxT = 1.3025, p = 0.51) or number of head-dipping events (Figure 2H; maxT = 0.853, p = 0.8). A significant treatment effect was found for time spent in SAP (Figure 2I; maxT = 2.5326, p = 0.04), but meaningful differences disappeared after adjusting *p*-values in *post-hoc* contrasts. Finally, a significant effect was found for SAP frequency (Figure 2J; maxT = 3.2229, p = 0.01), with flavone (p = 0.02) and diazepam (p = 0.01) reducing SAP.

**Figure 2:**
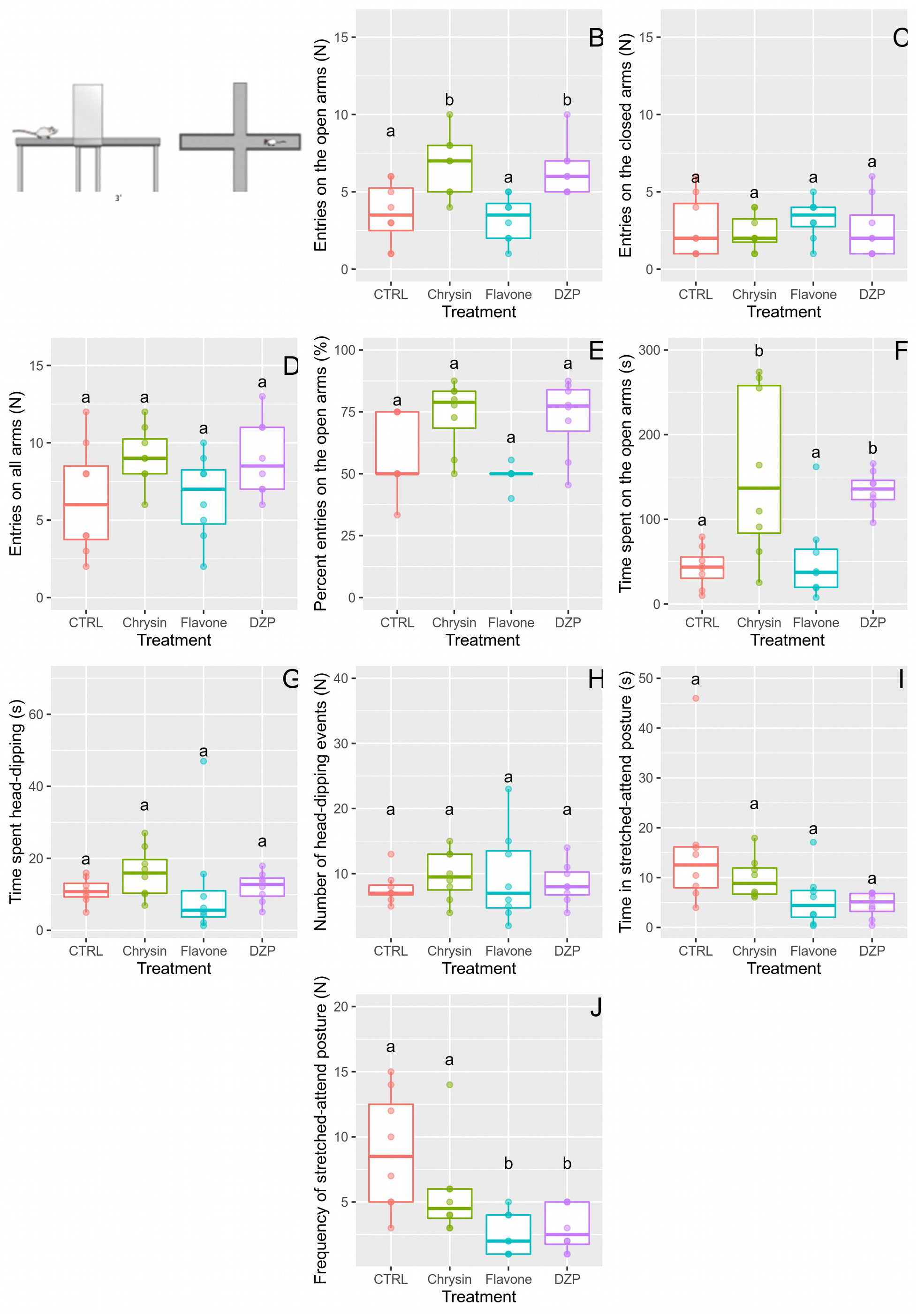
Effects of chrysin, flavone, and diazepam on the elevated plus-maze. A. Scheme of the apparatus. B. Entries on the open arms. C. Entries on the closed arms. D. Entries on all arms. E. Percent entries on the open arms. F. Time spent on the open arms. G. Time spent in headdipping. H. Number of head-dipping events. I. Time spent in stretched-attend posture (SAP). J. Frequency of SAP events. For variables with the same letter, the difference is not statistically significant. Likewise, for variables with a different letter, the difference is statistically significant (p≤0.05).

### Locomotor activity test

In the locomotor activity test (Figure 3A), none of the treatments modified the number of squares crossed (Figure 3B; maxT = 1.6296, p = 0.31) and grooming behavior (Figure 3C; maxT = 1.5029, p = 0.38); but significant differences were founded in rearing (Figure 3D; maxT = 4.0090, p = 0.01). *Post-hoc* test revealed that flavone backbone increased this variable (p = 0.01) respect to the control group.

**Figure 3:**
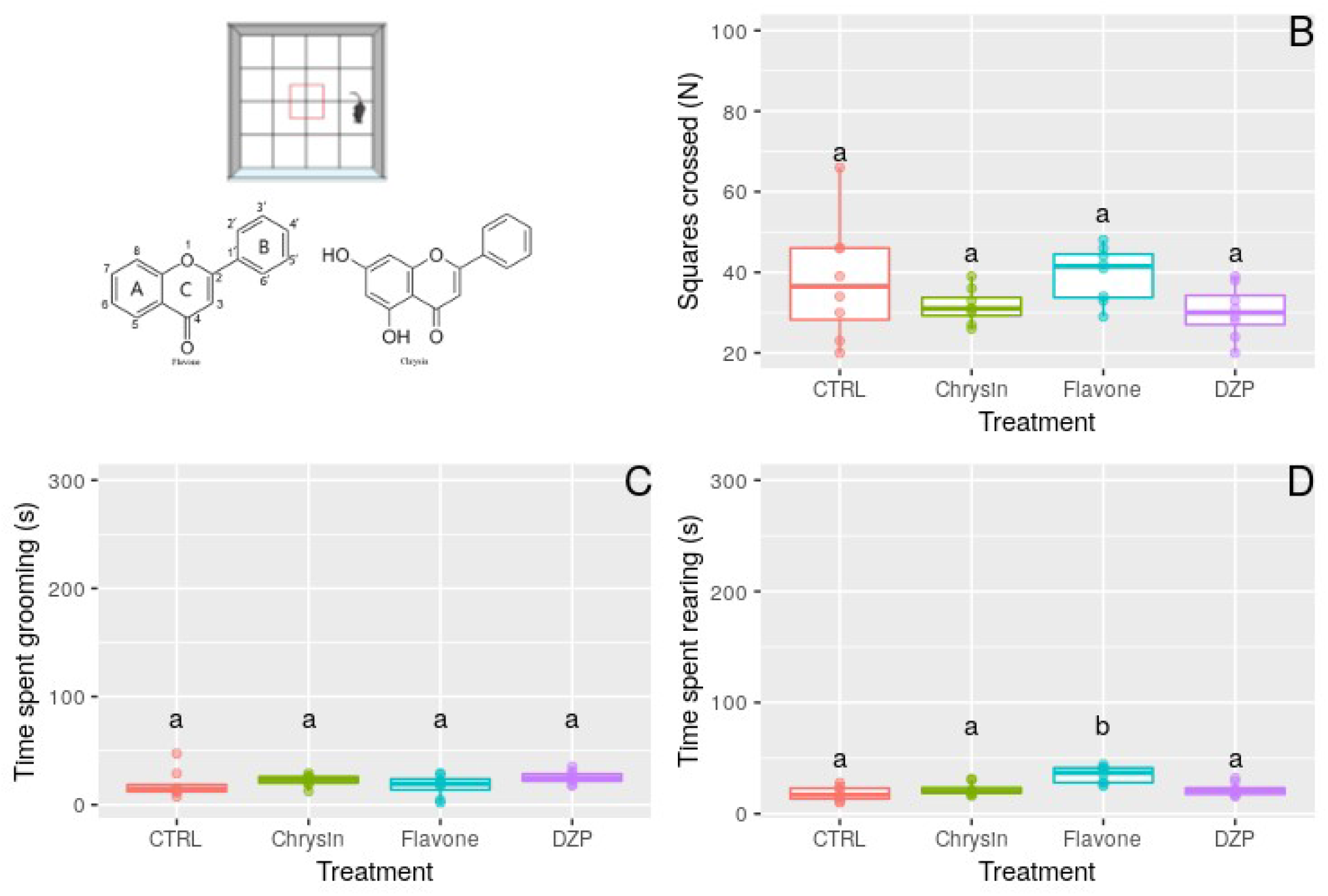
Effects of chrysin, flavone, and diazepam on the locomotor activity test. A. Scheme of the apparatus and test molecules. B. Squares crossed. C. Time spent in grooming. D. Time spent in rearing. For variables with the same letter, the difference is not statistically significant. Likewise, for variables with a different letter, the difference is statistically significant (p≤0.05).

### Light/dark test

In the light/dark test (Figure 4A), chrysin (p = 0.02) and diazepam (p = 0.02), but not the flavone backbone (p = 0.14), significantly increased (maxT = 3.7331, p = 0.01) the time spent into the white compartment (Figure 4B) compared to the control group. No significant changes were detected in the duration of entries (maxT = 2.0981, p = 0.12) or transitions (maxT = 0.6707, p = 0.89) to the white compartment (Figures 4C and 4D, respectively), associated with the treatments.

**Figure 4:**
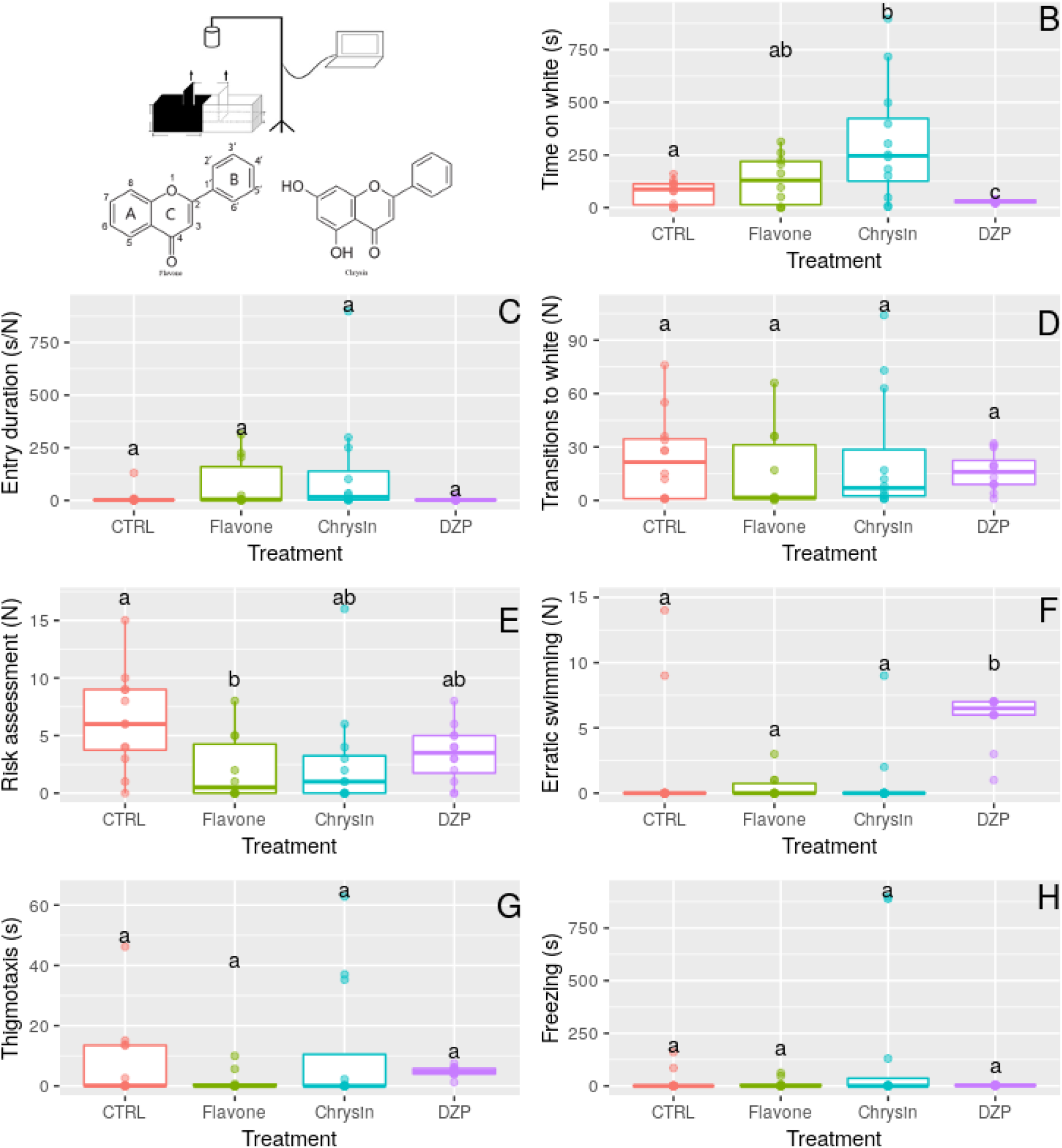
Effects of chrysin, flavone, and diazepam on the light/dark test. A. Scheme of the apparatus and test molecules. B. Time spent on the white compartment. C. Duration of entries on the white compartment. D. Transitions to white compartment. E. Frequency of risk assessment events. F. Frequency of erratic swimming events. G. Time spent in thigmotaxis. H. Duration of freezing events. For variables with the same letter, the difference is not statistically significant. Likewise, for variables with a different letter, the difference is statistically significant (p≤0.05).

On the other hand, the flavone backbone (p = 0.02), but neither chrysin (p = 0.06) nor diazepam (p = 0.06), significantly (maxT= 2.6404, p = 0.03) reduced risk assessment (Figure 4E) in relation to the control group. Diazepam (p = 0.031), but not chrysin (p = 0.60) or flavone (p = 0.50), increased erratic swimming (Figure 4F; maxT = 3.9468, p = 0.01). No significant changes were observed in thigmotaxis (Figure 4G; maxT = 1.4946, p = 0.39), or freezing duration (Figure 4H; maxT = 2.3935, p = 0.06) between treatments.

## Discussion

The present study demonstrated that 1) the anxiolytic-like effect of the flavonoid chrysin in the rat EPM and LDT, without locomotor effects, seems to be associated with hydroxyl groups substituted in 5 and 7 positions of the A ring in the flavone backbone; and 2) the flavone backbone is devoid of anxiolytic-like effects in the rat EPM in rats, but decreases risk assessment in the LDT in zebrafish and rearing in the locomotor activity test. These results contribute to the knowledge that positions of hydroxyl in the flavone backbone are important structural characteristic to produces its pharmacological action on the central nervous system, and suggest that a comparative approach is best suited to detect conserved drug effects across species that are more likely to be shared with humans.

In the present experiments, after chrysin treatment the number of entries into and time spent into the open arms were increased, without locomotor effects either in the EPM or in the locomotor activity test, an effect that is consistent with anxiolysis and is comparable to DZP. The effects of chrysin in the EPM have been shown before. Wolfman et al. (1994) showed that chrysin (1 mg/kg), isolated from *Passiflora coerulea*, exerts an anxiolytic-like effect in the EPM. We have previously shown that chrysin (2 and 4 mg/kg) blocked the anxiogenic-like effects of long-term ovariectomy in female rats, which were attributed to the activation of GABA_A_ receptors, as they were blocked by pretreatment with picrotoxin (Rodríguez-Landa et al., 2019).

Similar effects were observed in the LDT, a biobehavioral assay that is based in the preference of zebrafish for dark environments (Serra et al., 1999) and has been validated pharmacologically as a model to screening diverse anxiolytic drugs (Maximino et al., 2011; Magno et al., 2015). In this sense, the flavonoid chrysin increased the time spent in the white compartment in comparison to the control group. This variable is the main endpoint of the LDT (Maximino et al., 2011; Lima-Maximino et al., 2018), also clinically effective anxiolytic drugs that target different sites of the GABA_A_ receptor, like diazepam and other benzodiazepines, can increase it (Gebauer et al., 2011; Maximino et al., 2011; Magno et al., 2015), which has been considered as an anxiolytic-like effect at preclinical-research.

However, the flavone backbone did not modify the time spent into the open arms in the EPM neither the time spent in the compartment in the LD, in relation to the control groups. The basic flavone moiety (backbone) has a hydrogen bond donor site, lipophilic pockets, and an electron rich site (Marder & Paladini, 2002), which are thought to be important for the binding of both flavonoids and 1,4-benzodiazepines to the central benzodiazepine (BZD) site at GABA_A_ receptors (Marder & Paladini, 2002). Since the chrysin hydroxyl groups are located in one region of the flavone basic ring, this could be important for the steric interactions associated with binding, and therefore chrysin could bind to BZD sites with a higher affinity than flavone (Marder & Paladini, 2002; Huen et al., 2003). This could explain the wider anxiolytic-like effect of chrysin, but not flavone backbone, in the present study.

In addition to the effects of chrysin and flavone on the BZD site, another potential mechanism is a free radical scavenging effect: chrysin, like most flavonoids, exerts powerful antioxidant effects, because it possesses highly reactive hydroxyl groups in its basic structure, which confers a potent free radical scavenging activity (Panche et al., 2016). It has been reported that the generation of free radicals promote the inactivation of the enzyme glutamate decarboxylase (responsible for the synthesis of GABA), leading to a decrease of GABA concentrations and facilitating excitatory neurotransmission (Singh et al., 2014), and thus the scavenging potential of chyrsin could enhance the GABAergic neurotransmission (Harvey, 2015). On another hand, it has been reported that chrysin increases testosterone levels (Ciftc et al., 2011), which could be associated to its aromatase inhibitor activity (Campbell & Kurzer, 1993). In this sense, Fernández-Guasti et al. (2005) demonstrated that testosterone administration exerts an anxiolytic-like effect in male Wistar rats due to the GABA_A_-benzodiazepine receptor modulation by testosterone. This suggest that the flavonoid chrysin can exert more than one effect in the GABA_A_/benzodiazepine site complex.

In the other hand, chrysin did not alter risk assessment behavior in the present experiments, a variable that has been shown to be sensitive to non-benzodiazepine treatments as well as to benzodiazepines (Maximino et al., 2014; Lima-Maximino et al., 2018). In a multivariate approach, risk assessment has been shown to co-vary with other variables in the LDT, forming a separate cluster from thigmotaxis and time on white (Maximino et al., 2014); thus, risk assessment is sensitive to anxiolytic treatments, but represents a different dimension or factor in the multivariate structure of zebrafish behavior in the LDT.

Surprisingly, the flavone backbone decreased risk assessment in the LDT and SAP in the rat EPM, and increased rearing in the locomotor activity test. These outcomes are paradoxical; effects on rearing are difficult to interpret, but it has been suggested (Lever et al., 2006) that rearing is associated with environmental novelty, with a possible cognitive function in information-gathering for escape behavior. However, risk assessment in the LDT and SAP in the EPM can be interpreted as having an information-gathering function as well, suggesting different mechanisms for rearing, on the one hand, and risk assessment behavior in anxiety tests, on the other. Taken together, these results suggest that, although not able to reduce the avoidance of potentially adverse environments such as the open arm of the EPM or the light compartment of the LDT tank, the flavone backbone is able to reduce risk assessment behavior. Risk assessment has been suggested as a cognitive component of anxiety-like behavior (Blanchard et al., 2011), and is more sensitive than spatio-temporal measures (time spent on open arms) in the EPM to detect effects of serotonergic anxiolytics (Griebel et al., 1997).

Finally, the present study compared the effect of a single dose of chrysin (1 mg/kg) *versus* the flavone backbone. We have previously reported other doses (2 and 4 mg/kg) of this flavonoid with anxiolytic-like effect in the EPM (Rodríguez-Landa et al., 2019). However, that study used female Wistar rats with long-term ovariectomy, while in this work male Wistar rats were used. In other work, we demonstrated that, in male Wistar rats, only 1 mg/kg, but no 0.5 neither 2 mg/kg, of chrysin exerted an anxiolytic-like effect in the EPM in male Wistar rats (German-Ponciano et al., 2020). Thus, we used a single dose of chrysin in agreement with the ethical recommendations of the 3R’s principles in order to reduce the number of animals used in this study (Russell and Burch, 1959). Nevertheless, future work will be required in order to compare the possible dose-response curves of chrysin *versus* flavone backbone.

Overall the results of the present study show that, in zebrafish and rats, the anxiolytic-like effect of the flavonoid chrysin might be due to more than one effect in the GABA_A_/benzodiazepine receptor complex including its antioxidant effect, which could be related with the presence of the hydroxyl groups in its basic structure. However, additional work is required to determine the molecular mechanism through this nutraceutical compound and other flavonoids exert their potential therapeutic to ameliorate anxiety symptoms at preclinical research.

## Acknowledgments

This study was partially supported by SEP-CONACyT through funding from Programa de Fortalecimiento Académico del Posgrado de Alta Calidad (C-133/2014) to JFR-L. LJG-P received a fellowship from Consejo Nacional de Ciencia y Tecnología (CONACyT) for postgraduate studies in neuroethology (Reg. 297560) Additionally, he received a research scholarship to Brazil from Dirección General de la Unidad de Estudios de Posgrado de la Universidad Veracruzana.

## References

Bencan, Z., Sledge, D., & Levin, E. D. (2009). Buspirone, chlordiazepoxide and diazepam effects in a zebrafish model of anxiety. Pharmacology, Biochemistry and Behavior, 94, 75–80. https://doi.org/10.1016/j.pbb.2009.07.009

Blanchard, D. C., Griebel, G., Pobbe, R. L. H., & Blanchard, R. J. (2011). Risk assessment as an evolved threat detection and analysis process. Neuroscience & Biobehavioral Reviews, 35, 991–998. https://doi.org/10.1016/j.neubiorev.2010.10.016

Bortolotto, V. C., Pinheiro, F. C., Araujo, S. M., Poetini, M. R., Bertolazi, B. S., de Paula, M. T., Meichtry, L. B., de Almeida, F. P., de Freitas Couto, S., Jesse, C. R., & Prigol, M. (2018). Chrysin reverses the depressive-like behavior induced by hypothyroidism in female mice by regulating hippocampal serotonin and dopamine. European Journal of Pharmacology, 822, 78–84. https://doi.org/10.1016/j.ejphar.2018.01.017

Campbell, D. R., & Kurzer, M. S. (1993). Flavonoid inhibition of aromatase enzyme activity in human preadipocytes. The Journal of Steroid Biochemistry and Molecular Biology, 46, 381–388. https://doi.org/10.1016/0960-0760(93)90228-O

Chadha, R., Bhalla, Y., Nandan, A., Chadha, K., & Karan, M. (2017). Chrysin cocrystals: Characterization and evaluation. Journal of Pharmaceutical and Biomedical Analysis, 134, 361–371. https://doi.org/10.1016/j.jpba.2016.10.020

Ciftci, O., Ozdemir, I., Aydin, M., & Beytur, A. (2012). Beneficial effects of chrysin on the reproductive system of adult male rats. Andrologia, 44(3), 181–186. https://doi.org/10.1111/j.1439-0272.2010.01127.x

Cueto-Escobedo, J., Andrade-Soto, J., Lima-Maximino, M., Maximino, C., Hernández-López, F., & Rodríguez-Land, J. F. (2020). Involvement of GABAergic system in the antidepressant-like effects of chrysin (5, 7-dihydroxyflavone) in ovariectomized rats in the forced swim test: comparison with neurosteroids. Behavioural Brain Research, 112590. https://doi.org/10.1016/j.bbr.2020.112590

Diretriz brasileira para o cuidado e a utilização de animais para fins científicos e didáticos - DBCA. Anexo I. Peixes mantidos em instalações de instituições de ensino ou pesquisa científica, Resolução Normativa n. 30/2016 (2017).

Especificaciones técnicas para la producción, cuidado y uso de los animales de laboratorio, NOM (1999) (testimony of Norma Oficial Mexicana).

Fernández-Guasti, A., & Martínez-Mota, L. (2005). Anxiolytic-like actions of testosterone in the burying behavior test: role of androgen and GABA-benzodiazepine receptors. Psychoneuroendocrinology, 30(8), 762–770. https://doi.org/10.1016/j.psyneuen.2005.03.006

Filho, C. B., Jesse, C. R., Donato, F., Del Fabbro, L., de Gomes, M. G., Goes, A. T. R., Souza, L. C., Giacomeli, R., Antunes, M., Luchese, C., Roman, S. S., & Boeira, S. P. (2016). Neurochemical factors associated with the antidepressant-like effect of flavonoid chrysin in chronically stressed mice. European Journal of Pharmacology, 791, 284–296. https://doi.org/10.1016/j.ejphar.2016.09.005

Gambelunghe, C., Rossi, R., Sommavilla, M., Ferranti, C., Rossi, R., Ciculi, C., Gizzi, S., Micheletti, A., Rufini, S. (2003). Effects of Chrysin on Urinary Testosterone Levels in Human Males. Journal of Medicinal Food, 6, 387–390. https://doi.org/10.1089/109662003772519967

Gebauer, D. L., Pagnussat, N., Piato, Â. L., Schaefer, I. C., Bonan, C. D., & Lara, D. R. (2011). Effects of anxiolytics in zebrafish: Similarities and differences between benzodiazepines, buspirone and ethanol. Pharmacology, Biochemistry and Behavior, 99, 480–486. https://doi.org/10.1016/j.pbb.2011.04.021

Gerlai, R. (2014). Fish in behavior research: Unique tools with a great promise! Journal of Neuroscience Methods, 234, 54–58. https://doi.org/10.1016/j.jneumeth.2014.04.015

Germán-Ponciano, L. J., Puga-Olguín, A., Rovirosa-Hernández, M. D. J., Caba, M., Meza, E., & Rodríguez-Landa, J. F. (2020). Differential effects of acute and chronic treatment with the flavonoid chrysin on anxiety-like behavior and Fos immunoreactivity in the lateral septal nucleus in rats. Acta Pharmaceutica, 70(3), 387–397. https://doi.org/10.2478/acph-2020-0022

German-Ponciano, L. J., Rosas-Sánchez, G. U., Rivadeneyra-Domínguez, E., & Rodríguez-Landa, J. F. (2018). Advances in the preclinical study of some flavonoids as potential antidepressant agents. Scientifica, 2018, 1–14. https://doi.org/10.1155/2018/2963565

Ghosh, D., & Scheepens, A. (2009). Vascular action of polyphenols. Molecular Nutrition & Food Research, 53, 322–331. https://doi.org/10.1002/mnfr.200800182

Griebel, G., Rodgers, R. J., Perrault, G., & Sanger, D. J. (1997). Risk assessment behaviour: Evaluation of utility in the study of 5-HT-related drugs in the rat elevated plus-maze test. Pharmacology, Biochemistry and Behavior, 57, 817–827. https://doi.org/10.1016/S0091-3057(96)00402-9

Harvey, R. C. (2015). Benzodiazepines. In Small animal critical care medicine (pp. 864–866).

Hothorn, T., Hornik, K., van de Wiel, M. A., & Zeileis, A. (2006). A Lego system for conditional inference. The American Statician, 60, 257–263. https://doi.org/10.1198/000313006X118430

Huen, M. S., Hui, K. M., Leung, J. W., Sigel, E., Baur, R., Wong, J. T. F., & Xue, H. (2003). Naturally occurring 2′-hydroxyl-substituted flavonoids as high-affinity benzodiazepine site ligands. Biochemical Pharmacology, 66(12), 2397–2407. https://doi.org/10.1016/j.bcp.2003.08.016

Johnston, G. A. (2015). Flavonoid nutraceuticals and ionotropic receptors for the inhibitory neurotransmitter GABA. Neurochemistry International, 89, 120–125. https://doi.org/10.1016/j.neuint.2015.07.013

Kinkel, M. D., Eames, S. C., Philipson, L. H., & Prince, V. E. (2010). Intraperitoneal injection into adult zebrafish. Journal of Visualized Experiments, 42, 2126. https://doi.org/10.3791/2126

Krafczyk, N., Woyand, F., & Glomb, M. A. (2009). Structure-antioxidant relationship of flavonoids from fermented rooibos. Molecular Nutrition & Food Research, 53, 635–642. https://doi.org/10.1002/mnfr.200800117

Lawrence, C. (2007). The husbandry of zebrafish (*Danio rerio*): A review. Aquaculture, 269, 1–20. https://doi.org/10.1016/j.aquaculture.2007.04.077

Lever, C., Burton, S., & O’Keefe, J. (2006). Rearing on hind legs, environmental novelty, and the hippocampal formation. Reviews in the Neurosciences, 17, 111–133.

Leys, C., Ley, C., Klein, O., Bernard, P., & Licata, L. (2013). Detecting outliers: Do not use standard deviation around the mean, use absolute deviation around the median. Journal of Experimental Social Psychology, 49, 764–766. https://doi.org/10.1016/j.jesp.2013.03.013

Lima-Maximino, M. G., Cueto-Escobedo, J., Rodríguez-Landa, J. F., & Maximino, C. (2018). FGIN-1-27, an agonist at translocator protein 18 kDa (TSPO), produces anti-anxiety and anti-panic effects in non-mammalian models. Pharmacology Biochemistry and Behavior, 171, 66–73. https://doi.org/10.1016/j.pbb.2018.04.007

Magno, L. D. P., Fontes, A., Gonçalves, B. M. N., & Gouveia, A. (2015). Pharmacological study of the light/dark preference test in zebrafish (*Danio rerio*): Waterborne administration. Pharmacology, Biochemistry & Behavior, 135, 169–176. https://doi.org/10.1016/j.pbb.2015.05.014

Mangiafico, S.S. 2015. An R Companion for the Handbook of Biological Statistics, version 1.3.2.

Mani, R., Natesan, V., & Arumugam, R. (2017). Neuroprotective effect of chrysin on hyperammonemia mediated neuroinflammatory responses and altered expression of astrocytic protein in the hippocampus. Biomedicine & Pharmacotherapy, 88, 762–769. https://doi.org/10.1016/j.biopha.2017.01.081

Marder, M., & Paladini, A. (2002). GABA-A-Receptor ligands of flavonoid structure. Current Topics in Medicinal Chemistry, 2, 853–867. https://doi.org/10.2174/1568026023393462

Marventano, S., Godos, J., Platania, A., Galvano, F., Mistretta, A., & Grosso, G. (2018). Mediterranean diet adherence in the Mediterranean healthy eating, aging and lifestyle (MEAL) study cohort. International Journal of Food Sciences and Nutrition, 69(1), 100–107. https://doi.org/10.1080/09637486.2017.1332170

Maximino, C. (2018). Light/dark preference test for adult zebrafish (*Danio rerio*). Protocols.Io. https://doi.org/10.17504/protocols.io.srfed3n

Maximino, C., Brito, T. M. De, Dias, C. A. G. D. M., Gouveia Jr., A., & Morato, S. (2010b). Scototaxis as anxiety-like behavior in fish. Nature Protocols, 5, 209–216. https://doi.org/10.1038/nprot.2009.225

Maximino, C., da Silva, A. W. B., Araújo, J., Lima, M. G., Miranda, V., Puty, B., Benzecry, R., Picanço-Diniz, D. L. W., Gouveia Jr, A., Oliveira, K. R. M., & Herculano, A. M. (2014). Fingerprinting of psychoactive drugs in zebrafish anxiety-like behaviors. PLoS ONE, 9, e103943. https://doi.org/10.1371/journal.pone.0103943

Maximino, C., Lima-Maximino, M. G., & Siqueira-Silva, D. H. (2019). Zebrafish (*Danio rerio*) Environmental Summary, LaNeC (Marabá/PA, Brazil). Protocols.Io. https://doi.org/10.17504/protocols.io.rupd6vn

Maximino, C., Marques, T., Brito, D., Waneza, A., Manoel, A., Morato, S., & Gouveia, A. (2010a). Measuring anxiety in zebrafish: A critical review. Behavioural Brain Research, 214, 157–171. https://doi.org/10.1016/j.bbr.2010.05.031

Maximino, C., Silva, A. W. B. da, Gouveia Jr., A., & Herculano, A. M. (2011). Pharmacological analysis of zebrafish (*Danio rerio*) scototaxis. Progress in Neuro-Psychopharmacology & Biological Psychiatry, 35, 624–631. https://doi.org/10.1016/j.pnpbp.2011.01.006

National Research Council (US) Committee for the Update of the Guide for the Care and Use of Laboratory Animals. (2011). Guide for the Care and Use of Laboratory Animals (8th ed.). National Academy Press. https://www.ncbi.nlm.nih.gov/books/NBK54050/

Norton, W., & Bally-cuif, L. (2010). Adult zebrafish as a model organism for behavioural genetics. BMC Neuroscience, 11, 90. http://www.biomedcentral.com/1471-2202/11/90

Panche, A. N., Diwan, A. D., & Chandra, S. R. (2016). Flavonoids: An overview. Journal of Nutritional Science, 5, e47. https://doi.org/10.1017/jns.2016.41

R Core Team. (2019). R: A language and environment for statistical computing (3.4.4). http://www.r-project.org/

Ramanathan, V., & Thekkumalai, M. (2014). Role of chrysin on hepatic and renal activities of Nω-nitro-l-arginine-methylester induced hypertensive rats. International Journal of Nutrition, Pharmacology, Neurological Diseases, 4, 58–63. rcompanion.org/rcompanion/. (Pdf version: rcompanion.org/documents/RCompanionBioStatistics.pdf.)

Rodríguez-Landa, J. F., Hernández-López, F., Cueto-Escobedo, J., Herrera-Huerta, E. V., Rivadeneyra-Domínguez, E., Bernal-Morales, B., & Romero-Avendaño, E. (2019). Chrysin (5,7-dihydroxyflavone) exerts anxiolytic-like effects through GABA_A_ receptors in a surgical menopause model in rats. Biomedicine & Pharmacotherapy, 109, 2387–2395. https://doi.org/10.1016/j.biopha.2018.11.111

Russell, W. M. S., & Burch, R. L. (1959). The principles of humane experimental technique. Methuen.

Salgueiro, J. ., Ardenghi, P., Dias, M., Ferreira, M. B. ., Izquierdo, I., & Medina, J.. (1997). Anxiolytic natural and synthetic flavonoid ligands of the central benzodiazepine receptor have no effect on memory tasks in rats. Pharmacology, Biochemistry & Behavior, 58, 887–891. https://doi.org/10.1016/S0091-3057(97)00054-3

Sathiavelu, J., Senapathy, G. J., Devaraj, R., & Namasivayam, N. (2009). Hepatoprotective effect of chrysin on prooxidant-antioxidant status during ethanol-induced toxicity in female albino rats. Journal of Pharmacy and Pharmacology, 61, 809–817. https://doi.org/10.1211/jpp.61.06.0015

Schaefer, I. C., Siebel, A. M., Piato, A. L., Bonan, C. D., Vianna, M. R., & Lara, D. R. (2015). The side-by-side exploratory test. Behavioural Pharmacology, 26, 691–696. https://doi.org/10.1097/FBP.0000000000000145

Serra, E. L., Medalha, C. C., & Mattioli, R. (1999). Natural preference of zebrafish (*Danio rerio*) for a dark environment. Brazilian Journal Of Medical And Biological Research, 32, 1551–1553.

Sheela, C. G., & Augusti, K. T. (1995). Antiperoxide effects of S-allyl cysteine sulphoxide isolated from *Allium sativum* Linn and gugulipid in cholesterol diet fed rats. Indian Journal of Experimental Biology, 33, 337–341.

Siddiqui, A., Akhtar, J., Uddin M.S., S., Khan, M. I., Khalid, M., & Ahmad, M. (2018). A naturally occurring flavone (chrysin): Chemistry, occurrence, pharmacokinetic, toxicity, molecular targets and medicinal properties. Journal of Biologically Active Products from Nature, 8, 208–227. https://doi.org/10.1080/22311866.2018.1498750

Singh, P., Singh, D., & Goel, R. K. (2014). Phytoflavonoids: Antiepileptics for the future. International Journal of Pharmacy and Pharmaceutical Sciences, 6, 51–66.

Souza, L. C., Antunes, M. S., Filho, C. B., Del Fabbro, L., de Gomes, M. G., Goes, A. T. R., Donato, F., Prigol, M., Boeira, S. P., & Jesse, C. R. (2015). Flavonoid Chrysin prevents age-related cognitive decline via attenuation of oxidative stress and modulation of BDNF levels in aged mouse brain. Pharmacology, Biochemistry & Behavior, 134, 22–30. https://doi.org/10.1016/j.pbb.2015.04.010

Taiwo, A.E., Leite, F.B., Lucena, G.M., Barros, M., Silveira, D., Silva M.V., & Ferreira VM. (2012). Anxiolytic and antidepressant-like effects of Melissa officinalis (lemon balm) extract in rats: Influence of administration and gender. Indian Journal of Pharmacology, 44, 189–192. https://doi:10.4103/0253-7613.93846

Walf, A.A. and Frye, C.A. (2007). The use of the elevated plus maze as an assay of anxiety-related behavior in rodents. Nature Protocols, 2(2), 322–328, 2007. https://doi.org/10.1038/nprot.2007.44

Wolfman, C., Viola, H., Paladini, A., Dajas, F., & Medina, J. H. (1994). Possible anxiolytic effects of chrysin, a central benzodiazepine receptor ligand isolated from *Passiflora coerulea*. Pharmacology, Biochemistry & Behavior, 47, 1–4. https://doi.org/10.1016/0091-3057(94)90103-1

Yamamoto, Y. (2014). Effects of Dietary Chrysin Supplementation on Blood Pressure and Oxidative Status of Rats Fed a High-Fat High-Sucrose Diet. Food Science and Technology Research, 20, 295–300. https://doi.org/10.3136/fstr.20.295

Zanoli, P., Avallone, R., & Baraldi, M. (2000). Behavioral characterisation of the flavonoids apigenin and chrysin. Fitoterapia, 71, S117–S123. https://doi.org/10.1016/S0367-326X(00)00186-6

